# Platelet-specific P2Y_1_ receptor deficient mice have suppressed leukocyte recruitment in response to lipopolysaccharide

**DOI:** 10.1101/2024.11.04.621858

**Authors:** Dingxin Pan, Tolga Oralman, Reah Evans, Oliver Baker, Graham Cocks, Clive P. Page, Simon C. Pitchford

## Abstract

A role for the P2Y_1_ receptor in inflammation has been established using a pharmacological approach over an acute 4 hour time span. However, nucleotide-structure P2Y_1_ receptor antagonists have limited experimental use due to inadequate pharmacokinetics, and inability to decipher global versus cell specific effects *in vivo*. The creation of a conditional knock out (platelet) P2Y_1_ transgenic mouse model was designed to overcome these restrictions.

A homozygous P2Y_1_ *LoxP* mouse colony was created using CRISPR/Cas9 technology, and crossed with a hemizygous P2Y_1_ *LoxP* with PF4-cre to provide offspring that are homozygous for P2Y_1_ *LoxP* flanked allele, and hemizygous for the PF4cre (platelet P2Y_1_^-/-^) and offspring homozygous for P2Y_1_ *LoxP* flanked allele, but non-carriers for PF4cre (control mice). Animals were intranasally administered LPS to induce pulmonary inflammation to assess the influence of phenotype on leukocyte recruitment.

24 hours post intranasal LPS administration; pulmonary neutrophil and platelet recruitment were significantly suppressed, despite the fact that neutrophils retained the ability to migrate to fMLP *ex vivo*. Circulating platelet and leukocyte numbers were not different between control and platelet P2Y_1_^-/-^ animals. Tail bleeding times revealed the platelet P2Y_1_^-/-^ mice had a severe bleeding phenotype.

This is the first demonstration of a platelet specific P2Y_1_^-/-^ mouse model to confirm the importance of platelet P2Y_1_ receptors in the regulation of inflammatory responses with a 60-70% inhibition of leukocyte recruitment over an extended time period compared to previous pharmacological studies. platelet P2Y_1_^-/-^ mice will help further elucidate the mechanisms by which P2Y_1_ receptors regulate platelet activation during inflammation.

**Key Points:**

- Mice selectively deficient in the platelet P2Y_1_ have suppressed leukocyte and platelet recruitment during inflammation.
- We provide a methodology to determine mechanistic relevance that is otherwise limited by pharmacological approaches.

## Introduction

It is now recognised that platelets play a considerable role in the inflammatory response to infection, trauma, and inflammatory disorders.^1,2^ One important mechanism by which platelets participate in inflammation is via the intravascular interactions this cell type has with leukocytes and endothelium to efficiently orchestrate adhesion and migratory events involved in the leukocyte recruitment cascade.^3,4^ Studies in animals experimentally depleted of platelets with specific anti-platelet antibodies reveal leukocyte recruitment (as an indicator of inflammation) can be suppressed by over 80%.^5-11^ However such methodologies are challenging to further investigate mechanisms.^12^ Given the importance of the purinome in inflammation and haemostasis, we have previously investigated the role of platelet purinergic receptors in the regulation of inflammatory responses.^13-16^ The use of selective P2Y_1_, P2Y_12_, P2Y_14_ and P2X_1_ antagonists revealed a requirement for P2Y_1_ (and P2Y_14_) in both allergic and non-allergic models of pulmonary leukocyte recruitment instigated by exposure to allergen (in sensitised animals) or LPS that are also platelet-dependent.^17,18^ However, the use of antagonists poses difficulties in 1. Interpreting overall physiological relevance of any receptor (if the pharmacokinetic profile of the antagonist is not optimal), 2. confidence in the anatomical locality of where various cell types are being inhibited (due to tissue distribution), 3. a limited ability to adapt platelet depletion models to ascertain overall platelet input (due to the dynamics of antibody depletion).

As well as expression on platelets, it is notable that P2Y_1_ receptors are also expressed in CNS tissue and endothelial cells.^19^ We have previously developed platelet transfusion techniques in mice depleted of platelets long term using the bone marrow specific toxin busulfan, to elucidate the platelet-specific traits involved in leukocyte recruitment and the role of P2Y_1_ receptors.^17,20^ However, the methodology involved in such experimental approaches requires a high degree of technical skill and absolute precision of dosing regimens with busulfan to ensure the effect is selective for platelets and not affecting other blood elements. Furthermore, there is a need for *in vitro* incubation of platelets with compounds before reinfusion of this cell type to facilitate the interpretation of the data. Moreover, there are potential substantial animal welfare issues with long term cell depletion and reinfusion protocols. Various transgenic technologies have now been developed that provide platelet specific interventions.^21^ We have therefore developed a PF4 cre driven P2Y_1_ floxed murine model to further assess the physiological importance of platelet P2Y_1_ receptors in the regulation of pulmonary leukocyte recruitment in response to LPS.

## Materials and Methods

### Materials

ADP (Cat #01905), and the chemotactic peptide N-formylmethionyl-leucyl-phenylalanine (f-MLP) (Cat #F3506), prostaglandin E_1_ (PGE_1_) (Cat #P5515-1MG), LPS (from *Escherichia coli*, O55:B5 serotype), and urethane (Cat #U2500) were all purchased from Sigma Aldrich. The HTS Transwell 96-well plates (3μm pore size) (Cat #10077792) and RPMI 1640 cell media with L-glutamine (Cat #21875-034) were purchased from Thermo Fisher Scientific. Phycoerythrin (PE)-conjugated rat IgG (Cat #553930), Fluorescein isothiocyanate (FITC)-conjugated rat IgG (Cat #553988), and PE-anti-CD41 antibody (Cat #558040) were obtained from BD Biosciences. FITC-anti-P2Y_1_ antibody (Cat #APR-021-F) was purchased from Alomone Labs. Stromatol (Cat. #321200S) was purchased from Mascia Brunelli, Italy. Flow-Count Fluorospheres (beads, Cat #7547053) was from Beckman Coulter.

### Creation of conditional knock-out (cKO) of *P2ry1* in mice

The *P2ry1*^*flox/flox*^ (C57BL/6J-*P2ry1*^*em1Kcl*^) mouse was generated using CRISPR/Cas9 by the Genome Editing and Embryology Core facility at King’s College London, UK. All work was conducted in accordance with the UK Animal (Scientific Procedures) Act 1986, under project license PP9218930, with mice housed in individually ventilated cages and with a Specific Pathogen Free health status. To create the *P2ry1*^*flox/flox*^ line, loxp sites were positioned to flank exon 1 of *P2ry1* (ENSMUSE00000172824), containing the entire coding region, whilst ensuring that a lncRNA located on the reverse strand (ENSMUSG00000102564) remained intact prior to recombination (**Suppl Figures 1A and 1B**). To facilitate genotyping, a unique restriction site (BamHI) was inserted adjacent to each loxp site (**Suppl Figure 1B**). To reduce the likelihood of chromosomal re-arrangements during editing the 3’ and 5’ loxP sites were knocked-in sequentially (**Suppl Figures 1C**). In both cases a gRNA complexed to spCas9 was used along with a single-stranded oligonucleotide donor (ssODN) (**Suppl Figure 1B**). To generate the gRNAs, crRNA was annealed with tracrRNA followed by *in vitro* complexing with Cas9 protein and co-electroporated with ssODN donors (all from Intergated DNA Technologies) using a Nepa21 (Nepagene) into mouse zygotes (0.5dpc) from C57BL/6J mice (JAX® Mice Strain; Charles River).^22^

To generate the *P2ry1*^*flox/flox*^;Pf4^icre/wt^ line, an IVF was conducted with *P2ry1*^*flox/flox*^ female and *Pf4*^*icre/wt*^ male donors purchased from Jackson Labs (C57BL/6-Tg(Pf4-icre)Q3Rsko/J, strain 008535). *P2ry1*^*flox/wt*^;Pf4^icre/wt^ x *P2ry1*^*flox/flox*^ matings were then used to produce the required *P2ry1*^*flox/flox*^;Pf4^icre/wt^ line for experimental purposes.

### Mouse model of LPS-induced lung inflammation

All studies were carried out under the Animals (Scientific Procedures) Act of 1986 (United Kingdom) with local ethical approval of King’s College London. Litter group matched male and female WT or Plt-P2Y_1_^-/-^ mice were challenged with 1 mg/kg LPS in 50 μl via intranasal administration under isoflurane anaesthetic. 4 h and 24 hours post LPS challenge, 1.5 ml of bronchoalveolar lavage (BAL) fluid was collected and processed for total and differential cell counts as previously described.^18^ Circulating platelet and leukocyte numbers were enumerated at 4 h post LPS challenge after blood was taken via tail bleed, and quantified in stromatol (1:100 dilution) on an Improved Neubauer haemocytometer using an Axioskop Microscope under a ×40 objective.

### Measurement of bleeding time

Animals were kept under continuous anaesthesia with inhaled isoflurane anaesthetic. Bleeding assays were performed 4 h post LPS challenge in mice by tail tip amputation, immersing the tail in saline at 37 °C and continuously monitoring bleeding patterns as previously described.^18^ Each animal was monitored for up to 10 min and bleeding times determined using a stop clock. At the conclusion of the experiment, animals were killed with an overdose of urethane anaesthetic (25% w/v i.p.).

### Flow cytometry to measure P2Y_1_ receptor expression

Whole blood was obtained from terminally anaesthetized mice via cardiac puncture and centrifuged at 300 rcf for 3 min at room temperature to obtain PRP. PGE_1_ (2.5 μM) was added to prevent activation during staining. For each sample, 50 μL of PRP was stained with 1 μL of PE-anti-CD41 antibody (BD Pharmingen, 558040, 0.2 mg/mL) and 1 μL of FITC-anti-P2Y_1_ antibody (Alomone Labs, APR-021-F, 0.2 mg/mL) antibodies for 30 min at 4°C in the dark. After staining, cells were fixed with 150 μL of 2% paraformaldehyde. Platelets were identified based on size and CD41 positivity, and P2Y_1_ expression was assessed in the CD41+ gate. Samples were analyzed on a Beckman Coulter Cytoflex flow cytometer, recording 10,000 events per sample.

### Platelet and Neutrophil chemotaxis assays

Blood from mice was collected by means of cardiac puncture under terminal anaesthesia (25% w/v urethane, i.p) and washed platelets were isolated as described previously.^17^ Washed platelets (5□×□10^7^□mL^−1^) were treated with 2-mM CaCl_2_ before stimulation with vehicle (PBS) or ADP (100nM). Platelets, (80□μL) were then added to the top insert of the 96-well Transwell plate (3-µm pore size), with chemoattractant in the bottom well (0/30□nM fMLP in RPMI 1640 cell media). Following 90-min incubation at 37°C, media from the bottom chamber was stained with Stromatol (1:0.5) and platelets were quantified using an Improved Neubauer haemocytometer and a Leica DM 2000 LED microscope with a ×40 objective lens.

Bone marrow-derived neutrophils were also tested for migration toward fMLP (30 nM) using 3-μm pore-sized wells, as previously described.^23,24^ In brief, cells were resuspended at a concentration of 1 × 10^7^/ml in chemotaxis assay buffer (RPMI 1640 and 10% heat-inactivated fetal calf serum). The cell suspension was then transferred to the top insert of the

Transwell plate and the chemotaxis procedure carried out as described above for platelets. Following incubation for 60 min at 37°C, the number of neutrophils that migrated into the bottom chamber was determined by a total cell count combined with a differential cell stain (Diff Quick, Gamidor) to identify neutrophils.

### Statistical Analysis

Data are expressed as mean ± SEM. Quantification of cells via microscopy was conducted with the experimenter blinded to the sample identity. All other studies were quantified by machine (plate reader or flow cytometer). Chemotaxis data is normalised to a negative control to give a chemotactic index (CI) of fold mean of control values, due to baseline variations between donors. Groups are of equal size and are indicated in figure legends. Power calculations were undertaken to provide an estimation of the minimum sample size to detect difference between two means, dependent on intra-group variability of assays based on previous published data,^25-27^ or pilot data. Data were analysed using by means of 1-way ANOVA, followed by Dunnett’s multiple comparison post-test, or a T test. A *P* value of less than 0.05 was considered significant.

## Results

### Platelet P2Y_1_^-/-^ mice have an absence of P2Y_1_ receptor expression on platelets associated with a suppression of the inflammatory function of chemotaxis *in vitro*

PCR was conducted to examine *LoxP* and *Cre* expression on each litter mate. An example of identification of a test/platelet P2Y_1_^-/-^ mouse (litter mate 6.2a. Homozygous for P2Y_1_-*LoxP* flanked allele and hemizygous for PF4-*cre)* and control ‘wild type’ mouse (litter mate 6.2e. Homozygous for P2Y_1_-*LoxP* flanked allele, and a non carrier for PF4-*cre*) (**Figure 1A**). This effectively suppressed the expression of P2Y_1_ receptors on platelets as measured by flow cytometry (**Figure 1B**).

**Figure 1.**
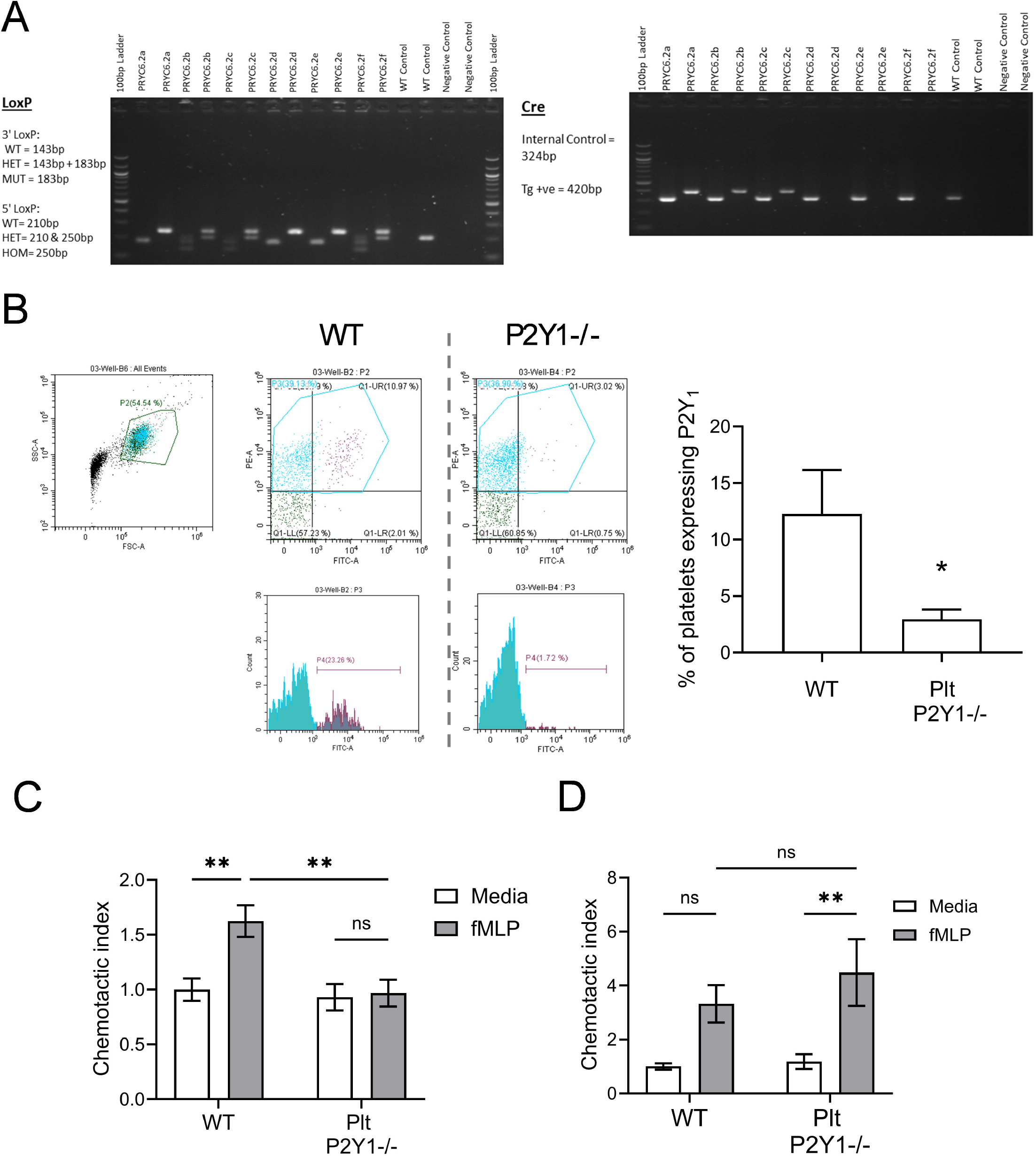
Characterization of platelets from C57BL/6J-*P2ry1*^*em1Kcl*^-Tg(Pf4-icre)Q3Rsko (Plt-P2Y_1_^-/-^ mice. Tissue taken from pups of PRYC x PRYP parents was used in PCR to determine offspring homozygous for P2Y_1_-*LoxP* flanked allele and hemizygous for PF4-*cre* (test mice) and offspring homozygous for P2Y_1_-*LoxP* flanked allele and a non carrier for PF4-*cre* (control ‘wild type’ mice). Representative PCR for *LoxP* and cre is shown of a litter (**A**). Platelets were taken via cardiac puncture and stained for with anti-CD41-PE conjugated antibody (platelet marker) and anti-P2Y_1_-FITC conjugated antibody to elucidate P2Y_1_ expression between ‘test’ (Plt-P2Y _1_ ^-/-^) and control (WT) platelets (**B**). In other experiments, platelets were harvested from blood, and leukocytes harvested from bone marrow of donor mice and their ability to migrate to fMLP (30nM) was measured using a transwell system (**C**) platelets, co-incubated with 100nM ADP, (**D**) neutrophils. Data expressed as means +/-SEM. n=3 (**B**) or 5-6 (**C**,**D**) per group. Significant difference represented: \**P* <0.05, \*\**P* <0.01.

As platelet activation via P2Y_1_ receptors is a necessary co-stimulus for their ability to migrate in response to inflammatory chemoattractants,^25,26^ we used a functional assay of chemotaxis to confirm that platelets harvested from Plt-P2Y_1_^-/-^ mice were unable to migrate to fMLP compared to platelets taken from WT mice (**Figure 1C**). For comparison, neutrophils isolated from the bone marrow of Plt-P2Y_1_ ^-/-^ mice were able to migrate in response to fMLP to the same degree as neutrophils harvested from WT mice (**Figure 1D**). Therefore, the P2Y_1_ receptor is confirmed to be essential for the ability of platelets to migrate toward an fMLP gradient.

### Platelet P2Y1^-/-^ mice have normal circulating levels of platelets and leukocytes, but exhibit an increased bleeding liability

We next investigated the effect P2Y_1_ receptor deficiency on platelets in a LPS-induced model of pulmonary inflammation. Plt-P2Y_1_^-/-^ mice had similar circulating levels of platelets (**Figure 2A**), neutrophils (**Figure 2B**) and mononuclear cells (**Figure 2C**) as WT mice. The intranasal administration of LPS (1mg/kg) at 4 hours, led to a decrease in circulating platelets in WT mice compared to mice administered saline, although this was not significant (**Figure 2A**). However, no effect on circulating platelet numbers was apparent in Plt-P2Y_1_^-/-^ mice (**Figure 2A**). There were also no differences in the circulating numbers of neutrophils and mononuclear cells between WT and Plt-P2Y_1_^-/-^ mice after LPS administration (**Figure 2B & 2C**). Thus, the haematopoietic phenotype of Plt-P2Y_1_ ^-/-^ mice was similar to WT mice based on these observations and functional data (**Figure 1D**). We also measured the effect of platelet P2Y_1_ deficiency on haemostasis through the measurement of tail bleeding time. As expected, Plt-P2Y_1_^-/-^ mice displayed a marked prolongation of bleeding compared to WT mice (**Figure 2D**), in both saline administered, and LPS-administered mice. It is also noted that inflammation did not affect bleeding, adding to evidence that the functions of platelets in inflammation are distinct to their participation in haemostasis.

**Figure 2.**
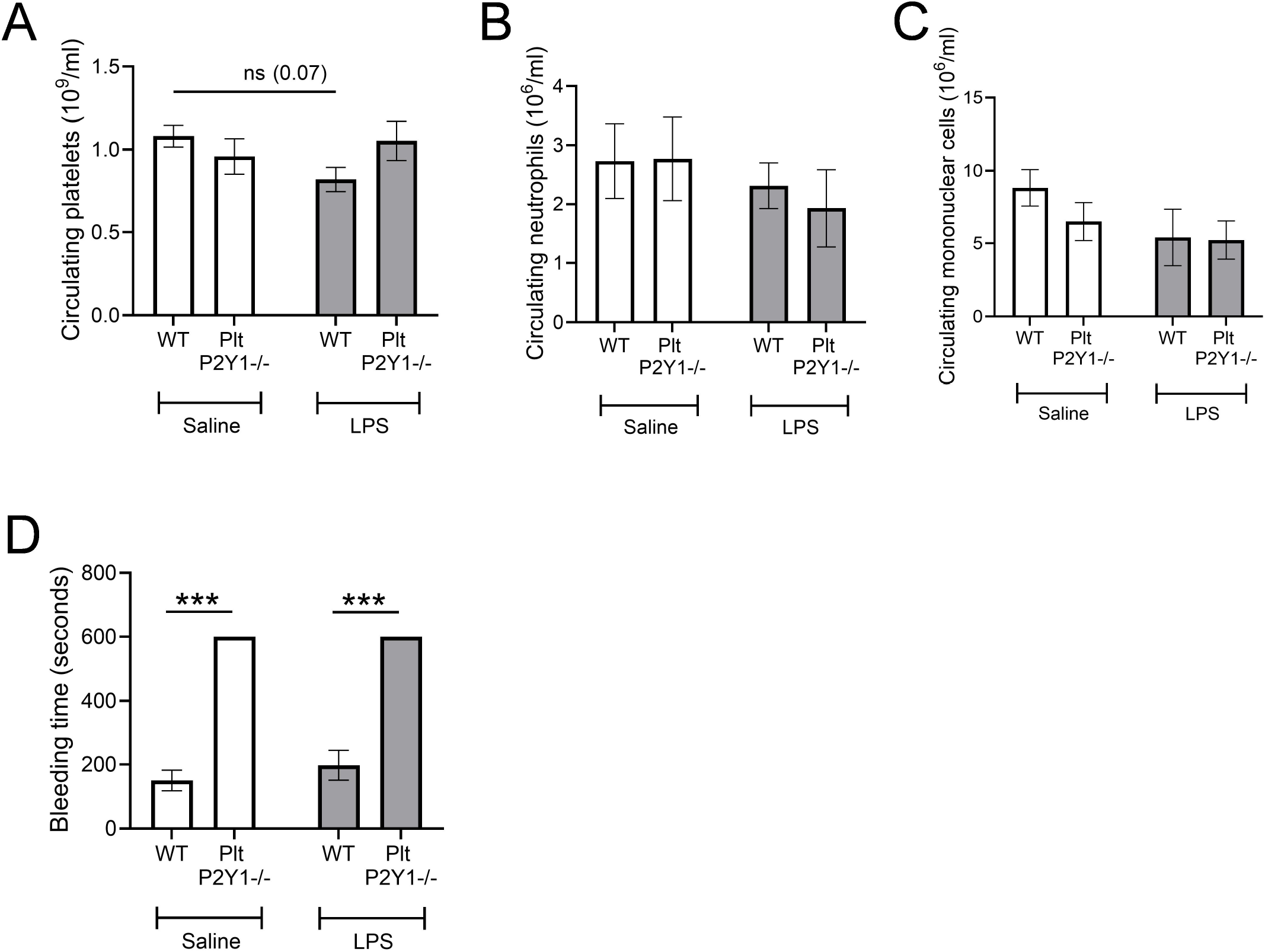
Measurement of haematological parameters in Plt-P2Y_1_^-/-^ mice. Mice were administered either saline or LPS (1mg/kg) intranasally and blood sample taken from tail vein after 4 hours and then anaesthetised with isofluorane. Blood was processed for enumeration of circulating platelets (**A**), neutrophils (**B**) and mononuclear cells (**C**). Time taken for tail veins bleeding to stop, maximum of 10 minutes (**D**). Data expressed as means +/-SEM. n = 5-6 per group. Significant difference: *** *P* <0.001.

### Mice with P2Y_1_ receptor deficient platelets have suppressed pulmonary leukocyte and platelet recruitment in response to LPS administration

LPS-induced pulmonary leukocyte recruitment in mice has been shown to be both platelet and P2Y1-dependent.^18,23^ In order to establish whether, and to what degree these dependencies are linked, pulmonary recruitment was quantified in Plt-P2Y_1_^-/-^ mice at 4 and 24 hours post LPS administration. Using equivalent levels of LPS to previous reports, this conditional (platelet) knock out mouse colony, despite being on a C57BL/6 genetic background, did not present with increased inflammatory cell recruitment at 4 hours post LPS administration (**Figure 3A-C**). However, at 24 hours, a robust inflammatory response was observed, with significantly increased pulmonary neutrophil and mononuclear cell recruitment compared to saline-treated control mice (**Figure 3D-F**). Plt-P2Y_1_^-/-^ mice were observed to have substantially reduced total cell counts (∼64% reduction **Figure 3D**), neutrophil counts (∼60% reduction **Figure 3E**), and mononuclear cell counts (∼100% reduction **Figure 3F**).

**Figure 3.**
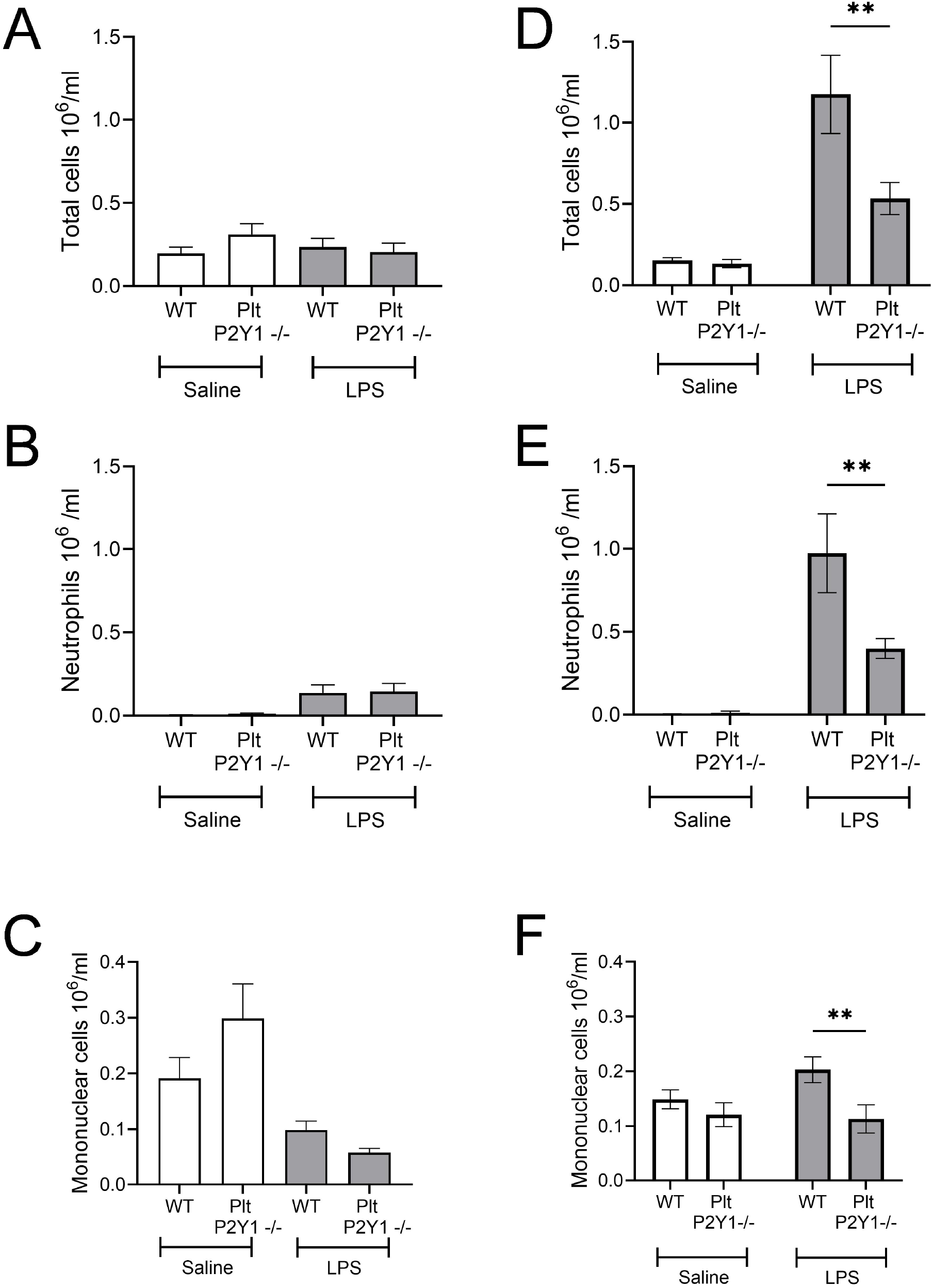
Measurement of pulmonary leukocyte recruitment in Plt-P2Y_1_ ^-/-^ mice. Mice were administered either saline or LPS (1mg/kg) intranasally and lavage samples undertaken at 4 hours (**A-C**) and 24 hours (**D-F**) post LPS, using terminal anaesthesia for enumeration of total leukocyte count (**A**,**D**); neutrophil (**B**,**E**) and mononuclear cell recruitment (**C**,**F**). Data expressed as means +/-SEM. n = 4 per group at 4 hours, and 7-8 per group at 24 hours. Significant difference: \*\**P* <0.01.

Localized platelet recruitment also occurs in response to intranasal LPS stimulation. This is not associated with the formation of pulmonary emboli, and whilst neutrophil independent, the mechanism of platelet recruitment is unknown.^27^ Flow cytometric analysis of lavage samples revealed an increased incidence of platelets after LPS administration, and this was significantly dependent on platelet P2Y_1_ receptors (**Figure 4A**). Analysis of lavage samples for the presence of red cells, revealed the increased platelet accumulation did not occur as a result of haemorrhage (**Figure 4B**). Furthermore, the deficiency of platelet P2Y_1_ receptors did not lead to increased haemorrhage in either saline or LPS treated groups (**Figure 4B**).

**Figure 4.**
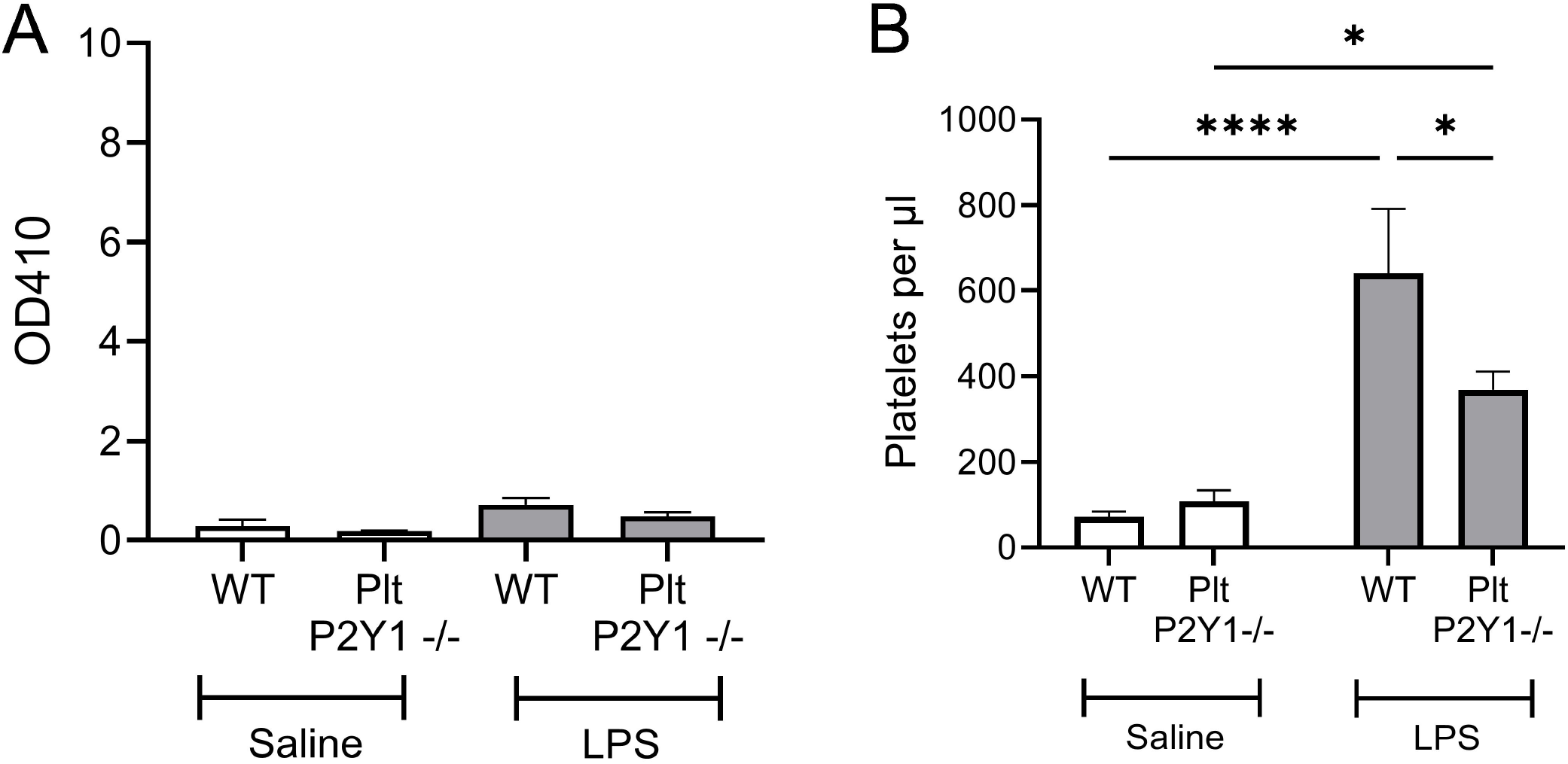
Measurement of pulmonary platelet recruitment and degree of haemorrhage in Plt-P2Y_1_^-/-^ mice. Mice were administered either saline or LPS (1mg/kg) intranasally and lavage samples undertaken at 24 hours post LPS, using terminal anaesthesia for measurement on platelet reader at 450nM of red cell contamination (**A**), and flow cytometric enumeration of platelets (**B**). Data expressed as means +/-SEM. n = 6-7 per group (**A**), and 13-15 per group (**B**). Significant difference: \**P* <0.05, \*\*\*\**P* <0.0001

## Discussion

This experimental approach confirms the importance of platelet activation via P2Y_1_ receptors in regulating pulmonary leukocyte recruitment after LPS administration. This adopted method of using conditional genetic knock-out technology to platelet expressed receptors has allowed us to distinguish the tissue localization of P2Y_1_ receptors in the regulation of leukocyte recruitment (i.e. allowing us to discount the expression of P2Y_1_ receptors on other inflammatory cell types, including endothelium, and on CNS tissues). Furthermore, we have developed a model that does not have associated methodological issues that require careful dose titration of platelet-ablative methods.^5,12^ The conditional knock-out of platelet P2Y_1_ receptors was made possible by the existence of PF4-*Cre* lines since PF4 is almost exclusively expressed by platelets and megakaryocytes.^28^ Despite recognition that the PF4 gene can be present in other haematopoietic stem cells, it is likely to be much lower than that seen in megakaryocytes.^21,29^ Other attempts to create platelet-specific *Cre* expression have used the GP1ba gene, which may affect platelet biology due to 1 intact copy, and imperfect *Cre* activation.^21^

The inflammatory response induced by administration of LPS in this model at 24 hours was substantial (around 1 million neutrophils/ml) and therefore provides a robust confirmation of the importance of P2Y_1_ receptors from previous reports that have used P2Y_1_ receptor antagonists for either neutrophil recruitment (around 0.3 million/ml) at 4 hours post LPS,^18^ or eosinophil recruitment (around 0.2-0.3 million/ml) after allergen exposure in previously sensitized mice.^17^ Whilst global PY_1_ deficient mice have not been studied in the same inflammatory models to understand the degree to which other P2Y_1_-expressing cell types might be involved, the inhibition of inflammation we have observed using a pharmacological approach (and therefore global, albeit brief due to the pharmacokinetic attributes of the drugs) in a similar model of LPS-induced inflammation was smaller (around 50%, at 4 hours with 3mg/kg of the selective P2Y_1_ antagonist MRS2500) to that observed here.^18^ Whilst pharmacological intervention with purinergic receptor antagonists that have a nucleotide structure can be effective over a longer period, they require multiple doses, and are effective with moderate rather than severe inflammatory responses.^17^ The LPS model has been used here as a proof of principle to show that platelet P2Y_1_ receptors are required for WBC recruitment *in vivo* during inflammation.

Therefore, the degree of suppression of leukocyte recruitment observed in this present study provides confidence for a major role for P2Y_1_ receptors in platelet activation involved in leukocyte recruitment. The extracellular ‘purinome’ is a critical feature of immune responses and inflammation, as it is for platelet haemostasis during blood clotting. However, these processes are distinct, and whilst we have been unable to reveal a role for platelet P2Y_12_ receptor or the ion channel P2X_1_, as others have;^30-32^ we have reported platelet activation by P2Y_14_ receptors in an inflammatory context.^18,33^ The production of this Plt-P2Y _1_ ^-/-^ mouse strain will therefore be a useful tool in understanding the interactions between different platelet-expressed purinergic receptors during the inflammatory response compared to their role in thrombosis and haemostasis. The functional consequences of activation of platelets via P2Y_1_ receptors in inflammation has been shown to require non-canonical RhoGTPase signalling involved in shape change, adhesion and motility (RhoA, Rac1), rather than the canonical PLC signalling pathway required for ADP-induced aggregation. Other endogenous nucleotides can activate platelets via P2Y_1_ receptors via RhoGTPase signalling to cause inflammatory-related functions in a biased manner,^26^ and so the use of Plt-P2Y_1_^-/-^ mice will help provide understanding as to the importance of this phenomenon *in vivo*.

In conclusion, the creation of a conditional knock out for platelet P2Y_1_ receptors confirms the major role for this purinergic receptor in regulating platelet-dependent leukocyte recruitment, and separately platelet recruitment in a murine model of inflammation.

## Funding Acknowledgement

This research was funded by a Medical Research Council (MRC) project grant [MR/T015845/1] awarded to Dr Simon Pitchford.

## Author Contributions

D.P., S.C.P, designed and performed the research, analyzed the data, and wrote the paper. O.B. designed the strategy to create the transgenic model. O.B. and G.C. managed, and T.O., and R.E. conducted procedures to create the transgenic model. C.P.P. co-wrote the paper. S.C.P. conceived the project, designed research, wrote the paper.

## Data Availability Statement

The data that support the findings of this study are available from the corresponding author upon reasonable request. Some data may not be made available because of intellectual property rights, privacy or ethical restrictions.

## Figure legends

**Suppl Figure 1. Creation of Platelet-P2Y1**^**-/-**^ **conditional knock out mouse**. Identification of transcript of interest from the Genome Reference Consortium Mouse Build 38 patch release 6 (GRCm38.p6) (**A**). Decision for location of LoxP insertion around P2Y_1_ gene on exon 1 (**B**). Breeding strategy for electrocorporation of 3’ LoxP and 5’ LoxP (**C**).

