## Supplementary Figure 1 for "Platelet-specific P2Y_1_ receptor deficient mice have suppressed leukocyte recruitment in response to lipopolysaccharide"

### Suppl Figure 1

**A**

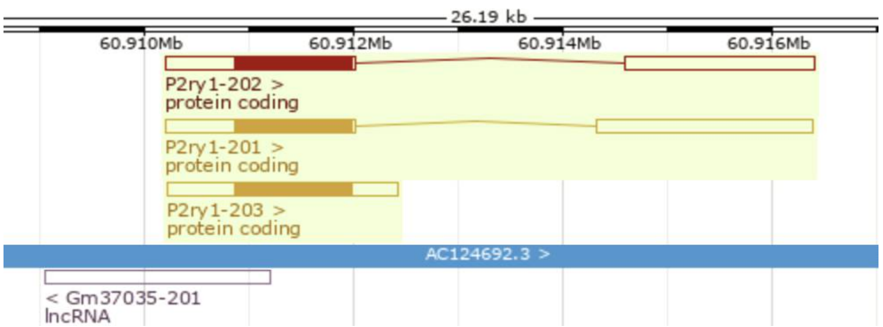


**B**

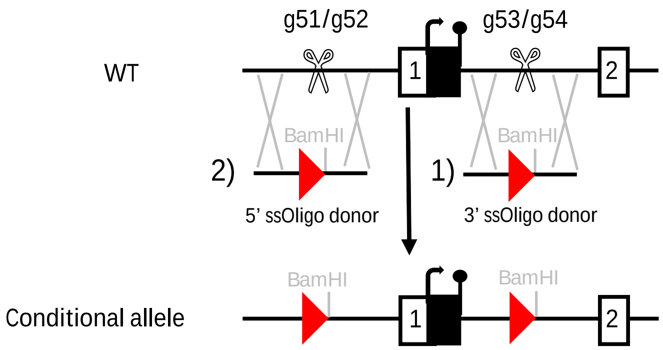


| gRNA | gRNA sequence 5' --> 3' | PAM |
|---|---|---|
| gRNA-51 | CAACTCCTTGAGTGTTTTTA | AGG |
| gRNA-52 | AACTCCTTGAGTGTTTTTAA | GGG |
| gRNA-53 | ATACTTCTATTCAGTAAGAA | AGG |
| gRNA-54 | TCAGTAAGAAAGGAGATTAC | AGG |

3’ ssOligo donor sequence (141mer): HA-loxP-BamHI-HA

5’- GAAGCATCAATTTGATGCACATATCCACATTTATTGTTCTTTCCTGTAATCTCCTATAACTTCGTATAGCATACATTATAC
GAAGTTATGGATCCTTCTTACTGAATAGAAGTAT
GTGTAATAAACACTTCCCCTTCCTCC - 3’

5’ ssOligo donor sequence (170mer): HA-loxP-BamHI-HA

5’- GGCTTGGCGGTAAGTGCCTAATGCCTTTAGTGGCTGAGACAGATCACCAACTCCTTGAGTGTTTTATAACTTCGTATAG
CATACATTATACGAAGTTATGGATCCTAAGGGAACAAA
GTCATTTAATATATGAAACTAGAATGAAATGGCTATAGAATATAATCGAAGGT 3’

**C**

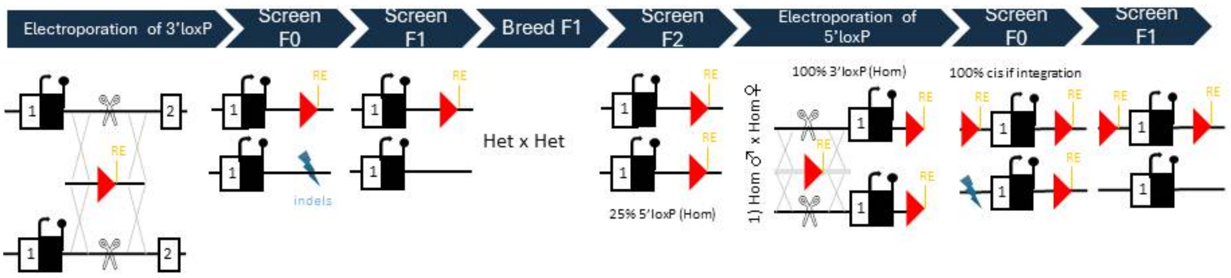


**Suppl Figure 1. Creation of Platelet-P2Y1$^{-/-}$ conditional knock out mouse.** Identification of transcript of interest from the Genome Reference Consortium Mouse Build 38 patch release 6 (GRCm38.p6) (**A**). Decision for location of LoxP insertion around P2Y$_1$ gene on exon 1 (**B**). Breeding strategy for electrocorporation of 3’ LoxP and 5’ LoxP (**C**).
